# Spatial structure, phase, and the contrast of natural images

**DOI:** 10.1101/2021.06.16.448761

**Authors:** Reuben Rideaux, Rebecca K. West, Thomas S. A. Wallis, Peter J. Bex, Jason B. Mattingley, William J. Harrison

## Abstract

The sensitivity of the human visual system is thought to be shaped by environmental statistics. A major endeavour in vision science, therefore, is to uncover the image statistics that predict perceptual and cognitive function. When searching for targets in natural images, for example, it has recently been proposed that target detection is inversely related to the spatial similarity of the target to its local background. We tested this hypothesis by measuring observers’ sensitivity to targets that were blended with natural image backgrounds. Targets were designed to have a spatial structure that was either similar or dissimilar to the background. Contrary to masking from similarity, we found that observers were most sensitive to targets that were most similar to their backgrounds. We hypothesised that a coincidence of phase-alignment between target and background results in a local contrast signal that facilitates detection when target-background similarity is high. We confirmed this prediction in a second experiment. Indeed, we show that, by solely manipulating the phase of a target relative to its background, the target can be rendered easily visible or undetectable. Our study thus reveals that, in addition to its structural similarity, the phase of the target relative to the background must be considered when predicting detection sensitivity in natural images.

## Introduction

The human visual system is tasked with parsing the complexity of natural environments into a coherent representation of behaviourally relevant information. These operations have been shaped by various selective pressures over evolutionary and developmental timescales. Therefore, the perceptual computations that guide cognition and behaviour ultimately serve to extract functional information from rich and complex naturalistic environments (Carandini et al., 2005; Field, 1987; Olshausen & Field, 2005; Parraga et al., 2000; Simoncelli & Olshausen, 2001). For example, a common task is to find a pre-defined target object in a complex or cluttered visual environment. The vast majority of our knowledge of the visual system, however, has been derived from experiments using relatively sparse stimulus displays that are not representative of our typical visual diets. The aim of the present study was to investigate how natural image structure influences target detection. We tested how detection is influenced by the spatial structure, phase, and contrast of natural image backgrounds to determine the features that best predict detection sensitivity.

Luminance contrast plays a critical role in most visual tasks. The human visual system is tuned to detect contrast across a range of spatial and temporal frequencies. Neurons in primary visual cortex (V1) are classically understood as processing local regions of oriented contrast that can define the borders of objects (Hubel & Wiesel, 1959). Such properties of individual neurons govern phenomenal perception and are thought to be shaped by the statistics of natural environments (Barlow, 1961, 1972). The encoding of contrast within the visual system is most commonly studied with oriented grating stimuli, such as Gabor wavelets. Grating stimuli are conveniently characterised by a simple set of parameters: orientation, contrast, position, and spatial frequency. From a computational perspective, “Gabor wavelet analyses” allow the decomposition of any image into mathematically tractable component features. Such analyses are relatively simple and are common in many computer vision applications. Early theory suggested that analogous decomposition processes occur in the visual system (Campbell & Robson, 1968). However, more recent studies suggest that individual visual neurons encode complex higher-order statistical information that is not necessarily predicted by Gabor parameters (e.g. Cadena et al., 2019).

One common approach to investigate contrast sensitivity in natural conditions is to have observers detect contrast-defined targets embedded in digital photographs or movies. Relative to sensitivity as typically quantified with a uniform background, spatio-temporal contrast sensitivity is diminished when viewing dynamic movies, particularly for lower frequencies (Bex et al., 2009). Furthermore, during free viewing of natural movies, the large-scale retinal changes caused by saccadic eye movements also diminish sensitivity likely due to forward and backward masking (Dorr & Bex, 2013; Wallis et al., 2015). In general, such studies have revealed that the sensitivity of the visual system does indeed depend on naturalistic context (Bex & Makous, 2002; Geisler, 2008).

Researchers have further sought to understand the statistical regularities of natural scenes that impact the detectability of targets. For example, various image structures, such as the density of edges within close proximity to the target, negatively impact detection sensitivity (Bex et al., 2009; see also Wallis et al., 2015). Indeed, the discriminability of visual objects can be predicted from the spatial proximity of surrounding visual clutter (Balas et al., 2009; Greenwood et al., 2010, 2012; Harrison & Bex, 2014, 2015, 2017; Rosenholtz et al., 2012; Wallis et al., 2019). More recently, it has been found that sensitivity scales inversely with the *structural similarity* between target and background (Sebastian et al., 2017, 2020). Structural similarity describes how similar two stimuli are in terms of the spatial distribution of phase-invariant contrast. Sebastian et al found that observers’ detection sensitivity decreases with increasing similarity. These studies thus predict that targets are most difficult to detect when they are similar to their backgrounds (Sebastian et al., 2017), particularly when those backgrounds are dense with edges (Bex et al., 2009). Other studies that have attempted to quantify the relationship between image statistics and sensitivity use post-hoc computational means to estimate the influence of natural image structure on target detection or apparent contrast (e.g. Haun & Peli, 2013; Wallis et al., 2015; Wallis & Bex, 2012). Very few studies, to the best of our knowledge, explicitly manipulated the consistency of a target’s appearance with the appearance of a natural image background in an experimental design (e.g. Neri, 2014, 2017; Teufel et al., 2018).

### The present study

The aim of the present study was to test observers’ sensitivity to targets presented on natural image backgrounds. Importantly, we designed the test stimuli a priori such that targets approximated the appearance of, and were aligned with, the local structure of a natural image background, or differed from the local structure. We therefore distinguish target-background *alignment* from target-background *similarity* in terms of the stimulus generation procedure (alignment) versus an image statistic (similarity). As shown in Figure 1, we automated the placement of targets within natural backgrounds according to oriented contrast energy at different image regions. We created two conditions, one in which targets were aligned with their backgrounds and one in which targets were misaligned with their backgrounds. In contrast to this stimulus generation procedure, target-background similarity is a measure of the correlation between a target and a background without a target. While target-background similarity ranges from 0 – 1 for all stimuli, our stimulus generation method results in higher similarity scores for aligned targets than misaligned targets (on average). Based on previous studies showing a negative impact of increasing target-background similarity on detection (Bex et al., 2009; Sebastian et al., 2017), we expected to find worse detection sensitivity when targets were aligned with the background – and were therefore highly similar – than when they were misaligned relative to the background – and were therefore relatively dissimilar. To anticipate our results, however, we found the opposite, instead revealing that the influence of target-background similarity on detection sensitivity depended almost entirely on the relative phase of the target.

**Figure 1.**
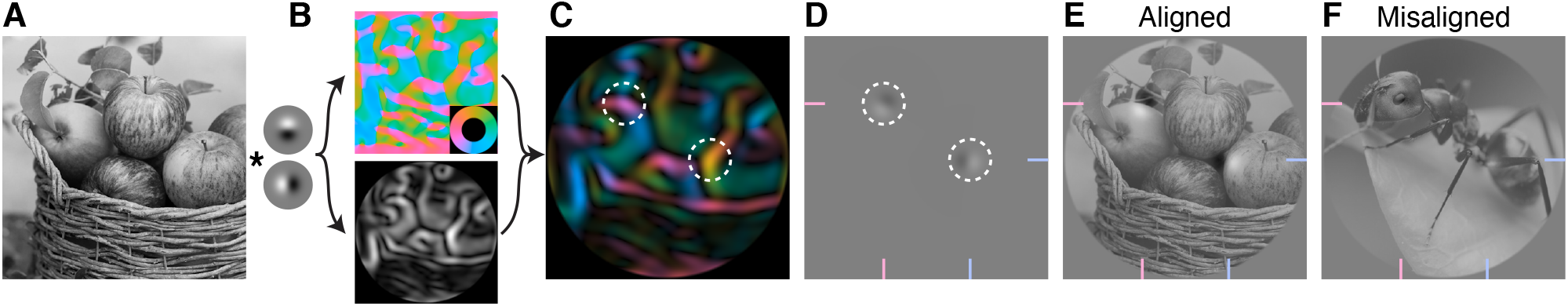
Stimulus generation method for testing sensitivity to contrast in natural images. A) An example source image taken from a collection of over 26,000 labelled images in the THINGS database (Hebart et al., 2019). B) We used a complement of derivative of Gaussian wavelets to filter each source image and compute the dominant orientation (top panel) and contrast energy (bottom panel) at each pixel location. Orientation is indicated by the inset colour wheel, which spans the full range (0 – 2pi) to indicate the preservation of the phase of the dominant filter. C) As shown by the white dashed circles, we selected target image regions according to the peaks of the oriented contrast maps. The number of targets varied from trial to trial from 1 to 16 in equally spaced log steps. D) Targets were oriented filters generated from the oriented contrast maps. These target features are thus aligned to the natural structure within the source image. Targets were then added to a natural image background, and observers were required to detect in which of two images the targets had been added. E and F) Targets were either aligned (E) or misaligned (F) with the background structure. Note that the same targets have been added to both examples but are more apparent in the aligned condition than the misaligned condition. As a guide, target filters are located at the intersection of pink and blue lines at the edges of panels D – F.

## Methods

### Participants

We used a single-subjects design in which we measured observers’ perceptual performance with high precision and treat each observer as a replication (Smith & Little, 2018). All observers were authors of the paper and had normal vision (RR, RW & WH). RW and RR were naïve to the specific experimental manipulations at the time of testing. The experiment was designed and carried out during a COVID-19 lockdown in Brisbane, Australia, in March 2021. Testing occurred, therefore, in each observer’s private residence.

### Design

We measured observers’ sensitivity to contrast changes in natural images in a 2 (target-background alignment: aligned or misaligned) x 5 (number of targets: 1, 2, 4, 8, or 16) x 5 (target amplitude: 0, 0.05, 0.1, 0.2, or 0.4 of maximum) design. Each observer completed 40 trials per condition for a total of 2000 trials in a fully within-participants design.

### Stimuli

Stimuli were programmed with the Psychophysics Toolbox (Brainard, 1997; Kleiner et al., 2007; Pelli, 1997) in MATLAB (v2018b, Mathworks) and were displayed either on a 15” MacBook Pro Retina or a 16” MacBook Pro Retina. Natural images of objects were taken from the THINGS database (Hebart et al., 2019), available via the Open Science Framework (https://osf.io/jum2f/). Images were converted to greyscale using the rgb2grey() function in MATLAB, and we assumed digital photos were encoded with a gamma of 2, and displays had a decoding gamma of 2.

We describe the stimulus generation process in detail below, but provide a brief overview here. On each trial, two different natural images were displayed, both of which were normalised in their contrast. Images had a diameter of 2° and were presented to the left and right of a red fixation spot. We chose this relatively small stimulus size for two reasons. First, we wanted to display images side by side, but close enough to central vision so as to mitigate effects of e.g. crowding. Second, because we were not able to monitor observers’ fixation compliance, the smaller stimulus size reduced the tendency for observers to make reflexive eye movements to high contrast image regions in their periphery. The target stimulus was the one in which wavelet filters had been blended with the natural image; the distractor was a natural image with no target filters. The filters were designed to be similar or dissimilar from the underlying natural image structure. We generated target stimuli by blending a source image of a natural object with derivative of Gaussian wavelets (henceforth: *filters*). The blending process followed four steps: 1) find the dominant orientation of each pixel in a source image, 2) find the relative contrast of each pixel in the image, 3) draw some number of filters at the highest contrast image regions, and 4) combine the filters with a source image. We expand on these steps below.

First, we used a steerable filter approach to determine the dominant orientation at every pixel in a given source image (Freeman & Adelson, 1991). Filters were directional first derivative of Gaussians oriented at 0° and 90°:

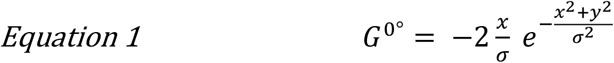

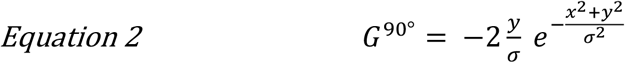

Where *σ* is the standard deviation of the Gaussian, *x* and *y* are the coordinates of each image pixel with point (0,0) at the centre, and G*^θ^* is the resulting filter. The Gaussian standard deviation was 0.08°. Within a trial, each filter was convolved with a source image:

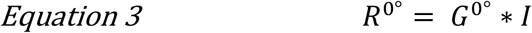

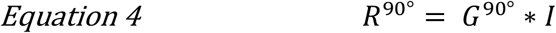

Where *I* is the source image with a mean of 0 and in the range [-1 1] and *R* is a filter response at each pixel location. We combined the filter responses to find the dominant orientation, 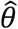, at each location:

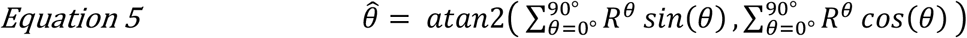

Second, we created a contrast map, *C*, of the filtered image by combining the filter outputs as follows:

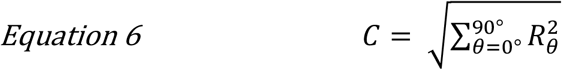

Third, we found the dominant orientation at the location of the contrast maxima:

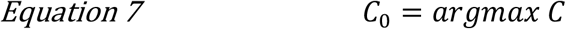

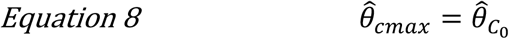

*C*_0_ indexes the x-y coordinates of the contrast maxima, and 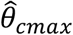 is the orientation at this location. We then steered a filter at this location as follows:

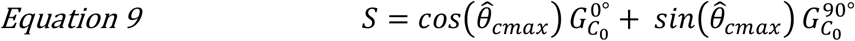

Here, *S* is the resulting filter stimulus in the range −1 to 1. Note the additional subscript of the filters that indicates that the filters were centred on the location of the contrast maxima, *C*_0_. This is trivially achieved by centring the x-y coordinates in Equations 1 and 2 on the coordinates of *C*_0_.

Finally, we created the target stimulus, τ, by combining the filter stimulus, *S*, with a normalised source image:

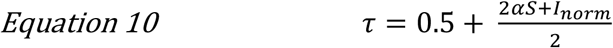

Where *α* is the filter amplitude expressed as a proportion of maximum possible contrast, and

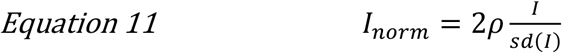

Here, *ρ* is the image root mean square (RMS) contrast, and *sd*(*I*) is the standard deviation of the source image. RMS contrast was set to 0.2 based on the findings of Sebastian et al (2017). The addition of 0.5 and denominator in Equation 10 normalises the range of the source image to [0 1] for display. Prior to this step, the target image was windowed in a circular aperture with a diameter matching the width of the source image (i.e. 2°) and a raised cosine edge, transitioning to zero contrast in 6 pixels. To constrain the filters generated by Equation 9 to appear within the windowed portion of the stimulus, the same aperture was applied to the contrast map, *C*, prior to generating the stimulus. Any values lower or higher than 0 or 1, respectively, in *I_norm_* were clipped.

For trials in which multiple filters were combined with a source image, we used an iterative procedure to draw *n* local maxima from the contrast image. Following the argmax operation in Equation 7, we updated the contrast map to minimise the contrast at the maxima:

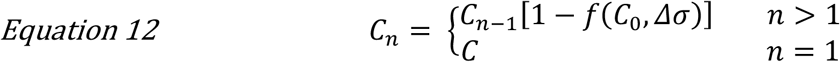

Where *C_n_* is the contrast map for the *n*-th filter, and *f*(*C*_0_,*Δσ*) is a two-dimensional Gaussian with a peak of 1 centred on the location of the maxima *C*_0_, and a standard deviation of *Δσ. σ* is the standard deviation of the basis filters, while *Δ* is a scaling factor that determines the spatial extent of change in the contrast map. The effect of this adjustment is the creation of a new local maxima at a different location than in the previous iteration. The greater the value of *Δ*, the greater the spatial spread of filters. The first filter location is always the image region with highest contrast. After accounting for the effect of *Δ*, subsequent filters are placed in regions of diminishing contrast. In trials in which multiple filters were present, backgrounds were randomly selected as described above.

### Image selection

The 26,107 images in the THINGS database are grouped into 1,854 concepts (e.g. “dog”, “cup”, “brush” etc), such that there are at least 12 unique, high quality images for each concept (Hebart et al., 2019). In each testing session (500 trials), we selected 1000 source images from unique concepts such that no two images were drawn from the same concept. The target background was thus always drawn from a different concept than the distractor image. However, it was necessary that some concepts were repeated across testing sessions, and it was also possible that some individual images were also repeated across sessions (but never within sessions).

On each trial, we selected two images from the set of 500: one image for the target background, and a second image was the distractor. On half the trials, the target filters were generated from the target background and were therefore aligned with the background, while on the other half of trials they were generated from the distractor image – but blended with the target background – and were therefore misaligned relative to the target background. Target filters were generated from the distractor background on misaligned trials, as opposed to an unused image, so that the filters and their source image were presented on every trial, but we doubt this decision was important to our results. We chose to present two different background images on each trial, rather than, for example, presenting two of the same background images, because we did not want observers to attempt to simply spot the difference between two similar images. Instead, observers had to perform a more natural task of searching unfamiliar and unique backgrounds for targets.

### Procedure

A typical trial sequence is shown in Figure 2. Each trial began with a small red fixation spot in the centre of the display, followed by outlines of the upcoming stimulus locations. Natural image backgrounds were followed by a blank of 500ms, after which time the observer reported which of the two patches contained the target filter(s) using the keyboard. Following the observer’s response, the image patch with target filters was re-displayed for an additional 500ms, outlined in green or red depending on whether the observer’s response was correct or incorrect, respectively. Feedback was provided to facilitate observers reaching a stable level of performance. No breaks were programmed but could be taken by withholding a response. Each session included ten repeats of each trial type, all presented in random order, giving 500 trials per session and taking approximately 15 minutes when no breaks were taken.

**Figure 2.**
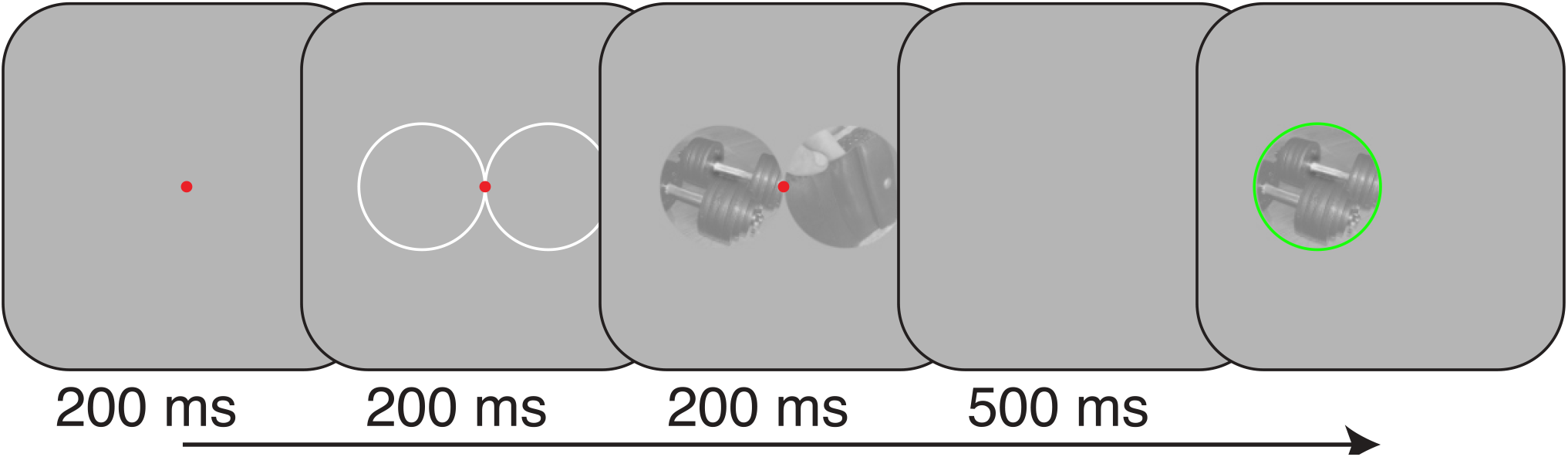
Schematic of typical trial sequence. The target and background could appear on the left or right of the screen with equal probability. Following an observer’s response, the background containing the target filters was framed by a green or red circle depending on whether the response was correct or incorrect, respectively.

### Sensitivity analyses

We quantified observers’ sensitivity to the target filters in natural images in a generalised linear model (GLM) framework. We describe the most pertinent aspects of the framework below, but, for the impatient reader, we note that these equations accumulate to the fitglme() function in MATLAB, or, equivalently, the lmer() function in R with the lme4 package (Bates et al., 2015).

In a standard single-interval detection paradigm in which a target is either present or absent, sensitivity, *d*’, is calculated as:

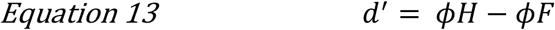

Where *ϕ* is the normal integral function, *H* is the proportion of hits, and *F* is the proportion of false alarms, under the assumption of equal variance. An observer’s criterion, *c*, is calculated as:

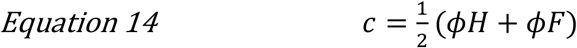

In a GLM, *d*’ and *c* (bias) are computed as predictor weights *β*_1_ and *β*_0_, respectively, that are passed through a probit link function, which is the normal integral function:

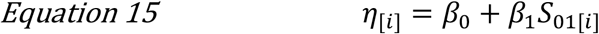

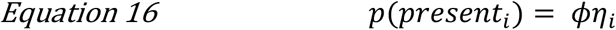

Where *η* is the sum of weighted linear predictors and *S*_01_ is the absence or presence of the signal (i.e. 0 or 1, respectively) on the *i-*th trial. By fitting such a probit model, estimates of the predictor weights *β*_1_ and *β*_0_ are identical to *d*’ and *c*, respectively, as calculated in Equation 13 and Equation 14. Whereas these equations fully specify sensitivity and bias in a single interval present/absent judgement task, some small modifications are needed to quantify sensitivity in a two-alternative forced choice task (2AFC) as in our experiment. First, *S*_01_ denotes whether the target filter(s) appeared in the left or right spatial interval, defined as -.5 or .5, respectively. Similarly, observers’ reports (i.e. “target appeared in the left or right interval”), were defined as −1 and 1, respectively. Finally, in a 2AFC, observers have two opportunities to detect the target – once per spatial interval – and so raw *d*’ will be greater than in a single-interval detection design. Therefore, sensitivity (but not bias) must be scaled by 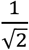 (Macmillan & Creelman, 2004):

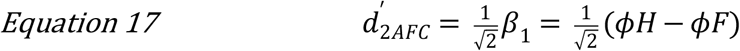

Importantly, we can extend Equation 15 to quantify sensitivity to any number of other predictors, *x_ω_*:

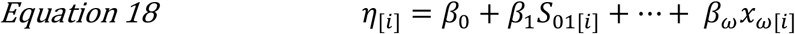

Consider, for example, the influence of filter amplitude (*α*) on an observer’s sensitivity:

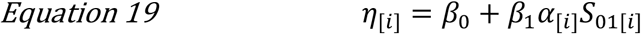

Note that filter amplitude is entered into the model as an interaction with target location because the model’s predicted outcome is a spatial report; target amplitude alone can only predict a change in bias. In preliminary model fits, we found that such bias was not significantly different from zero, and thus included only interactive terms to facilitate interpretability of the standard bias term, *β*_0_. We selected other model predictors according to the model that produced the lowest Akaike information criterion (AIC; see below).

Finally, we implemented this model as a multilevel GLM (GLMM) to partially pool coefficient estimates across observers (Gelman & Hill, 2007). By using a GLMM, we model each observer’s predictor weights as having come from a population distribution with mean *μ* and variance *σ*^2^:

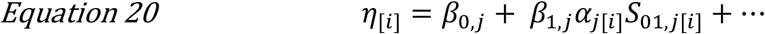

Where

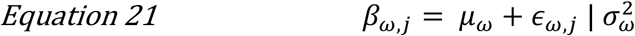

Here, *ϵ_ω,j_* is the offset for each predictor *ω* and observer *j*, relative to the parameter’s mean *μ*, contingent on the parameters’ estimated population variance *σ*^2^. The partial pooling of observers’ data in a GLMM results in more extreme values being pulled toward the population mean estimate. Note that in our experiment, however, such pooling is relatively minor due to the large number of trials, and therefore high precision, of each observer’s estimated performance, as well as the relatively small number of observers. Because images were drawn randomly from trial to trial from a pool of tens of thousands of images, we did not expect many, if any, repeats of each image. We therefore did not model the background images as a random effect, but we note that such a design could be chosen in future to estimate the variance associated with each tested background.

We entered into the model the factors target amplitude, number of filters, and target-background alignment, which, as noted above, were each entered as an interaction with the spatial interval of the target. In hindsight, our inclusion of the condition in which target amplitude was 0 was unnecessary. For all such trials, therefore, we set all predictors to have a value of 0 so they were omitted from model calculations. The model fit was improved by including nonlinear terms by raising amplitude and number of Gabors to the exponents 0.5 and 2, respectively. We further tested all combinations of interactions, but none improved the model fit as assessed by the Akaike information criterion.

### Post-hoc analysis of interaction between the number of filters and filter amplitude

We modelled the joint influence of number of filters and filter amplitude on proportion correct as a two-dimensional surface (see Figure 5B). The surface is defined as:

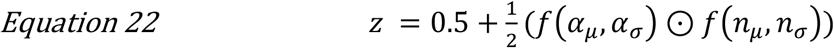

Where z is the surface of estimated proportion correct, f(α_μ_,α_σ_) is a cumulative probability function relating target amplitude to accuracy according to a threshold and variance, α_μ_ and α_σ_, respectively, and f(n_μ_,n_σ_) is a cumulative probability function relating the number of filters to accuracy according to a threshold and variance, n_μ_ and n_σ_, respectively. Here, ⊙ refers to the element wise product of cumulative distributions. This function can be thought of as a two-dimensional psychometric function, with separable means and standard deviations. The input parameters into the cumulative functions were free parameters, fit by minimising the summed squared error between the average proportion correct and *z* using Matlab’s fminsearch(). While there are no doubt other ways of quantifying the interaction between filter amplitude, number of filters, and proportion correct, this model suffices for our purposes.

### Structural similarity

We quantified the similarity between targets and their backgrounds using the same approach as Sebastian et al (2017 see their equation S9):

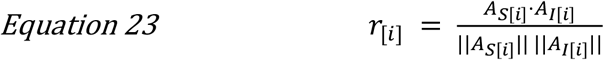

Where r_[i]_ is the similarity between the steered filter targets (i.e. *S* in Equation 9) and the background image *I* on trial *i*. A_s[i]_ is the Fourier amplitude spectrum of the filters, and A_I[i]_ is the Fourier amplitude of the natural image background, both of which are vectors and were computed by taking the absolute of the complex values of the Fourier transforms. r_[i]_ is thus a phase-invariant metric. Prior to computing Λ_I[i]_, we windowed the natural image background to include only the same regions as the locations of target filters. This was achieved by first computing a contrast map in which two-dimensional Gaussians were positioned at each target location. Gaussians had the same standard deviation as the target filters and had their peaks normalised to one. We then computed the elementwise product of the source image and this contrast map, which produced the background image entered into Equation 23. Note that our method to generate stimuli and target-background similarity both depend on oriented contrast within the frequency band of the target. Variations in structural similarity for the aligned and misaligned targets are shown in Figure 6.

## Results

We tested observers’ ability to detect target filters that were blended with natural image backgrounds. Targets were designed such that they were either aligned or misaligned with the structure of the background. We tested detection of 1, 2, 4, 8 and 16 target filters and across a range of target amplitudes. We first describe our results in terms of raw accuracy, and then report the results of our modelling analysis in which we quantified observers’ sensitivity in terms of *d*’.

### The influence of target amplitude, number of filters, and target-background alignment

The proportion of correct responses as a function of each factor is shown in Figure 3. The amplitude of the target most clearly impacts accuracy, such that accuracy increases approximately linearly with (log) target amplitude (Figure 3A). Although the relationship between proportion correct and the number of filters is less consistent (Figure 3B), there is a general increase in accuracy with increasing number of target filters. As described below, however, the relationship between number of filters and sensitivity was not significant. We also found a highly consistent effect of target-background alignment across observers (Figure 3C). Contrary to our expectations based on recent reports of similarity masking (Sebastian et al., 2017), however, we found better performance when target filters were aligned with their backgrounds than when they were misaligned with their backgrounds. This difference in performance is significant at the group level (*t*(2) = 5.862, *p* = 0.028, *d* = 3.38), but we more formally quantify the relationship between these factors in a GLMM below. Importantly, given the high measurement precision of these data (1000 observations per data point shown in Figure 3C), we can treat each observer as an independent replication of the effect, regardless of any specific inferential statistic (Smith & Little, 2018).

**Figure 3.**
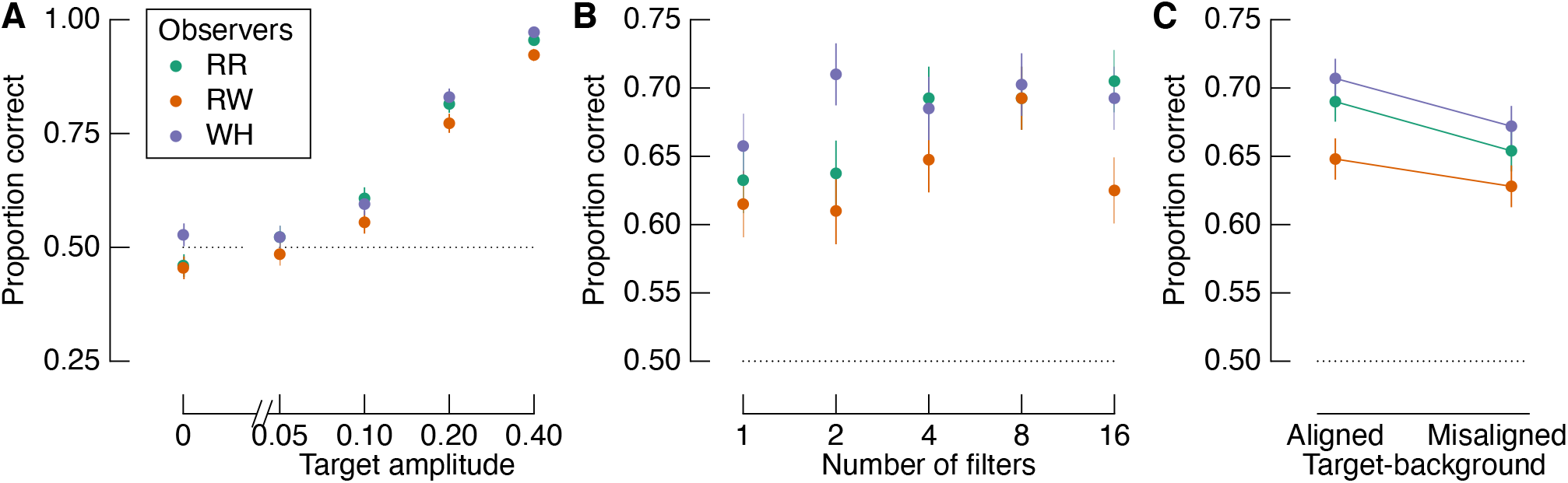
Proportion correct target identifications based on the three experimental factors: A) Target amplitude, B) Number of filters, and C) Target-background alignment. Colours represent different observers, as indicated by the legend in panel (A). The dotted line in each panel shows chance performance. There were 400 trials per data point in panels (A) and (B), and 1000 trials per data point in panel (C). Error bars show one binomial standard deviation.

We estimated detection sensitivity as a function of the experimental factors with a GLMM. Modelled sensitivity is shown in Figure 4 in the same format as Figure 3. Target amplitude and target-background alignment were both significant contributors to the model. Importantly, as shown in Figure 4C, sensitivity was greater when target filters were aligned with the background than when they were misaligned. The number of filters did not predict sensitivity, which is consistent with the relatively noisy relationship between accuracy and the number of filters as shown in Figure 3B.

**Figure 4.**
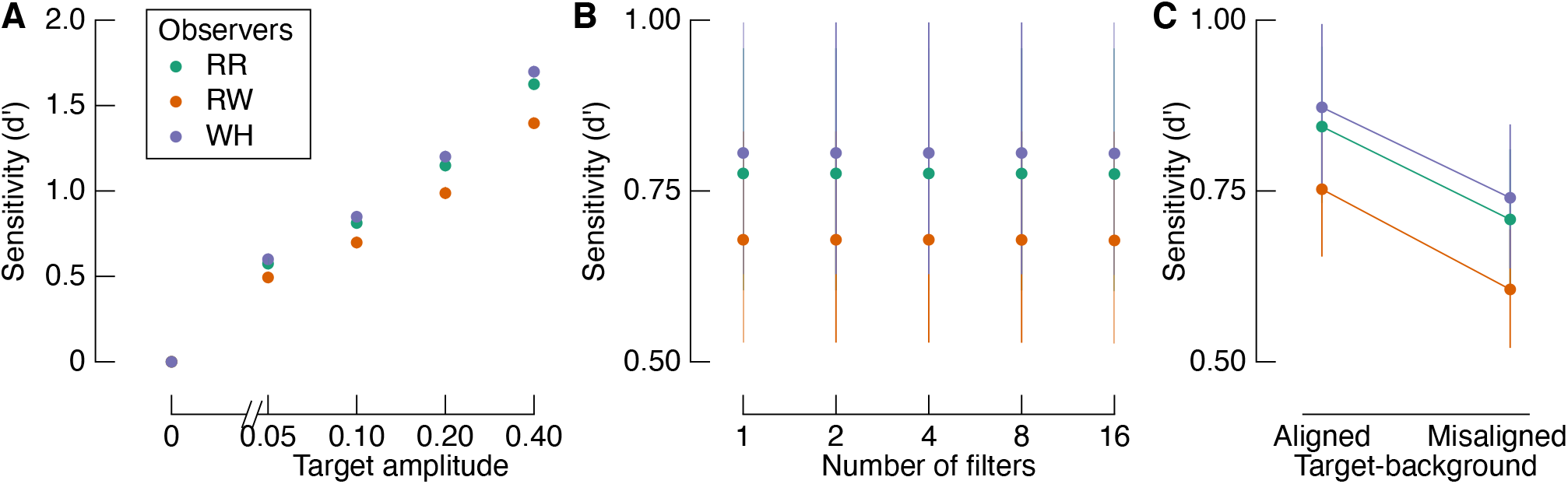
Modelled target detection sensitivity. A) Sensitivity varies systematically with target amplitude, but not the number of filters (B). C) Sensitivity depends on the target-background alignment, such that it is greater when filters are aligned with the background than when they are misaligned. Error bars in all panels show one standard error across marginalised conditions, but are smaller than the point size in (A).

We manipulated the number of filters because we expected to find an improvement in performance with increasing filter number. The lack of a main effect of filter number in the results above, therefore, was unexpected. Although not critical to our primary interest in the influence of target-background alignment, we tested whether the number of filters interacted with target amplitude using a more direct test than the full GLMM above. Data points in Figure 5A show mean proportion correct responses, marginalised to show the influence of the number of filters for each target amplitude. At the two highest target amplitudes tested (top two lines), there is indeed an effect of the number of filters: as the number of filters increases, so too does observers’ accuracy. The same data points are shown in Figure 5B arranged as a surface that maps proportion correct as conditional on the combination of conditions. We interpolated these points as a two-dimensional surface function that quantifies the interaction between number of filters and amplitude (see Methods). The warm colours clustering in the top right corner reveal that increasing the number of filters in the targets has the strongest effect at higher target amplitudes.

**Figure 5.**
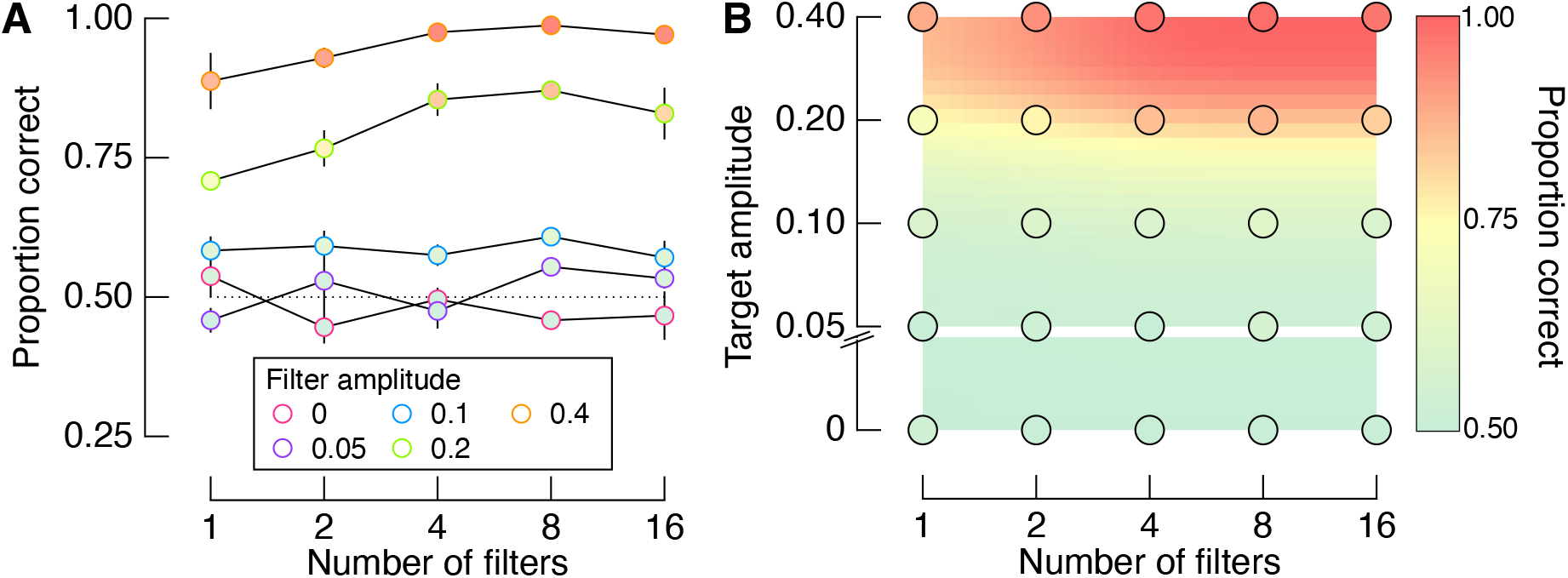
Interaction between the number of filters and filter amplitude. A) Proportion correct for each number of filters at each amplitude level, averaged across the three observers. B) Points show the same data as in (A), but arranged as a surface, while the background is interpolated from these points. Error bars in (A) show one standard error across observers, which is smaller than the point size in many cases.

### The influence of target-background similarity

The perceptual performance described above reveals that observers were better able to detect targets on natural image backgrounds when the targets were aligned with the underlying spatial structure of the background than when the targets were misaligned with the background. These results are contrary to our expectations based on the data of Sebastian et al (2017) who found that sensitivity is negatively correlated with the *structural similarity* between target and background, a metric that scales from zero (no similarity) to one (perfect similarity). We therefore next tested whether observers’ performance was instead *positively* correlated with target-background similarity using the same analysis of similarity as in this previous study (Equation 23).

Shown in Figure 6A are the histograms of similarity for all trials across observers, which, by design, can be separated according to the filter alignment relative to the background. The dashed vertical line shows the median similarity of all trials, regardless of target-background alignment. By isolating trials from each condition according to whether they fall above or below this arbitrary cut-off, we can test whether accuracy depends more on similarity or target-background alignment. Figure 6B shows the proportion of correct target detections for aligned and misaligned targets that were most and least similar to the background, respectively. As per the main analyses above, accuracy was greater for aligned than misaligned trials. The more diagnostic analysis is shown in Figure 6C, in which the proportion of correct target detections are shown for aligned trials that were *less* similar than the included misaligned trials (i.e. we limit the analyses to the tails of the similarity distributions). We again find that accuracy was higher for the aligned condition than the misaligned condition, despite the aligned targets having lower similarity with the background than the misaligned trials. Therefore, target-background alignment predicts performance much more strongly than target-background similarity.

**Figure 6.**
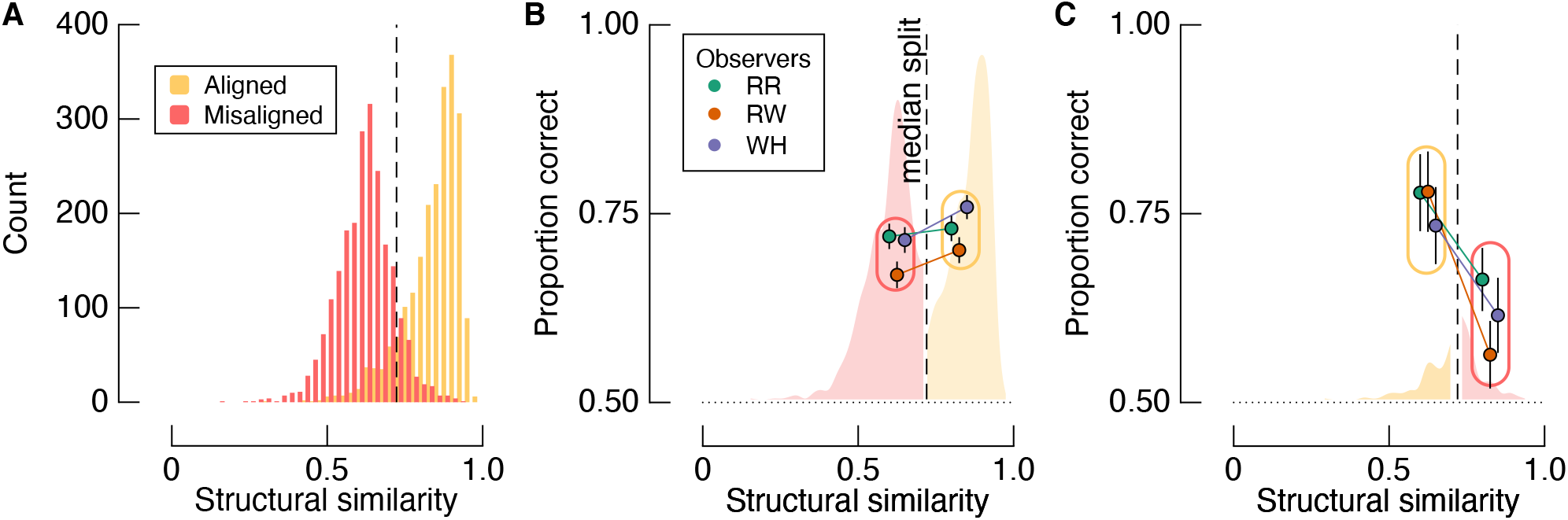
Analysis of the influence of target-background similarity on perceptual performance. A) Histograms show structural similarity between targets and background separately for the aligned and misaligned conditions. The vertical dashed line is the median similarity of all trials. B) Proportion correct for misaligned trials that were lower in similarity than aligned trials, and vice versa, as determined by median split. Accuracy for the misaligned trials is shown to the left of the median, and accuracy for the aligned trials is shown to the right of the median. C) Proportion correct for aligned trials that were lower in similarity than misaligned trials, and vice versa, as determined by median split. Accuracy for the aligned trials is shown to the left of the median, and accuracy for the misaligned trials is shown to the right of the median. Note that proportion correct is higher in the aligned condition regardless of similarity. Error bars in (B) and (C) show one binomial standard deviation.

In a final analysis of perceptual performance, we attempted to replicate the findings of Sebastian et al (2017) using the condition in our experiment that is most analogous to the one in theirs, namely, the misaligned condition. In both this previous study and the misaligned condition of the present study, the blending of target filters and their backgrounds did not depend on any structural alignment. Instead, any incidental alignment can be quantified in terms of structural similarity. We therefore tested whether we found an inverse relationship between target-background similarity and detection accuracy for the misaligned condition. Figure 7 shows proportion correct for trials in similarity bins (bin width = 0.1) for each observer. We modelled these data with logistic regression, with random intercept and slopes grouped by observer (i.e. a logistic GLMM). Fits are shown as solid lines in Figure 7. There is a clear negative relationship between structural similarity and accuracy, with a mean slope of −0.96 (population standard deviation = 0.75). Similarity was not a significant predictor in the model (*p* = 0.13), but the trends are nonetheless consistent across observers and also with the data of Sebastian et al.

**Figure 7.**
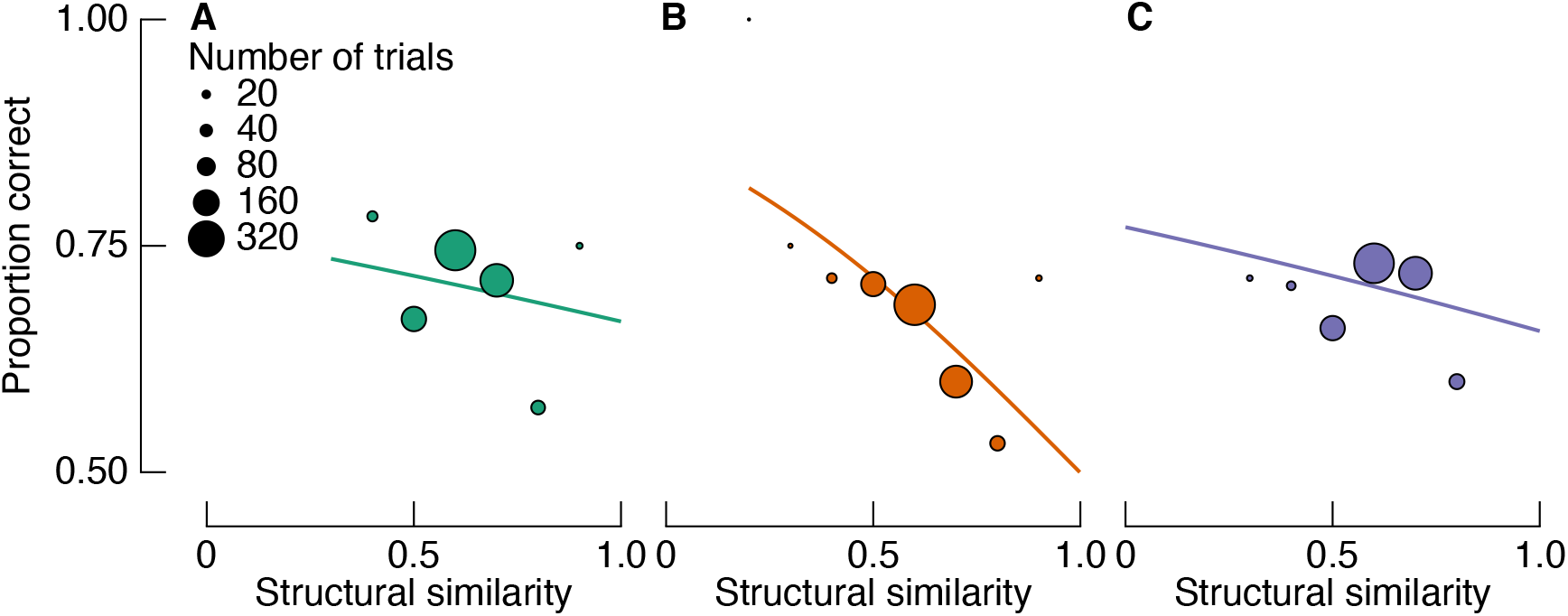
Proportion correct as a function of target-background similarity for misaligned trials only. A-C) show data for each observer, colour coded as in Figure 3. Solid lines are fits from a logistic GLMM, showing a negative relationship between similarity and detection accuracy, as reported by Sebastian et al (2017).

### Discussion

We found that observers are better at detecting contrast-defined targets when they are aligned with a natural image background compared with when they are misaligned with the natural image background. The superior performance on aligned than misaligned trials did not depend on the structural similarity of targets relative to backgrounds, in contrast to the results of Sebastian et al (2017). Because the target filters tended to be aligned with object edges (i.e. the points of highest contrast in natural images; see Figure 1 and Methods), these data also appear to contradict the findings of Bex et al (2009; see also Wallis & Bex, 2012). Bex et al found that sensitivity was lower in image regions of relatively high edge density, whereas we found higher sensitivity when targets were aligned with edges than when they are randomly positioned.

A potentially simple explanation for the discrepancy between our data and earlier work concerns the phase alignment of target filters and backgrounds: the *phase* of a target filter relative to the phase of the local natural image background determines local contrast. This fact is demonstrated in Figure 8A. When target filters are designed to be aligned with their background structure (left panels), local contrast depends strongly on phase. When the targets and background are phase-matched, target-background amplitude is additive, resulting in greater local contrast (top left panel). By inverting the phase of those same filters, target-background amplitude is subtractive, reducing local contrast (bottom left panel). Indeed, Bex and Makous (2002) speculated that this dependence of local contrast on phase alignment explains a loss of sensitivity to phase-scrambled natural images. We tested this hypothesis directly in Experiment 2. Note, however, that, while phase-mismatched target filters reduce local contrast, they may not be less visible. See, for example, the demonstrations in Figure 8B. While the phase-mismatched filter has a lower local contrast than the phase-matched filter, the phase-mismatched filter is conspicuous. Indeed, within each half of these images, the absolute change of luminance is the same, regardless of filter phase. It therefore remains an open question as to how this manipulation will affect observers’ sensitivity.

**Figure 8.**
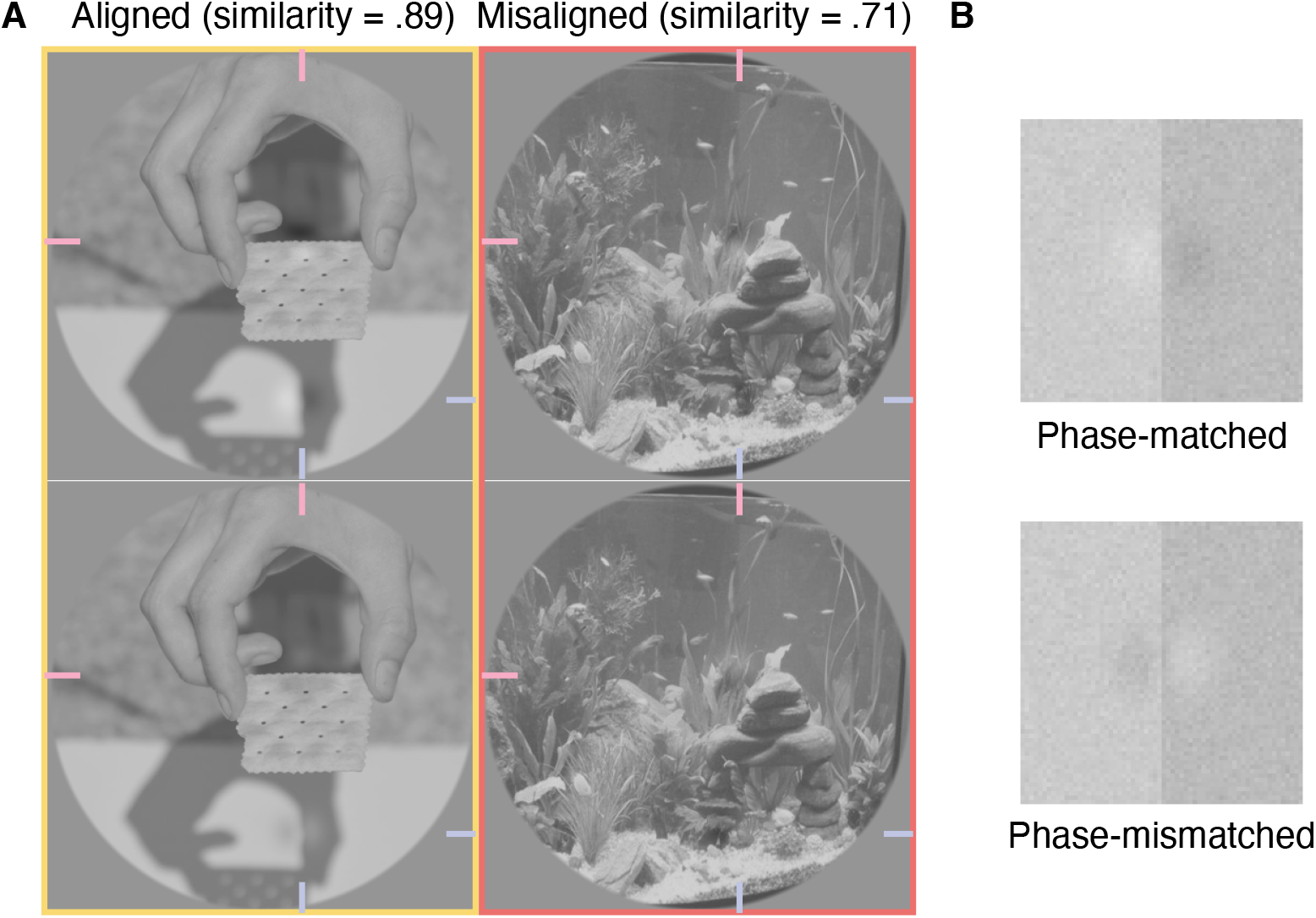
A demonstration of the influence of filter phase on local contrast, and examples of conditions tested in Experiment 2. A) Added to all panels are the same two target filters that were designed to be phase-matched with the local spatial structure in the top left panel. Filters in the bottom row have an inverse phase, and are therefore mismatched relative to the source image. Filters are most apparent in the top left panel: one filter is aligned with the top horizontal edge of the cracker, and the other is aligned to the left side of the vertical shadow of the thumb. Note that the structural similarity metric is phase invariant, and therefore the filters have the same similarity score within each column. Target filters are located at the intersection of pink and blue lines at the edges of each panel. In Experiment 2, we averaged observers’ performance over the misaligned conditions (right column), because phase alignment is relative only to the source image. B) Simplified demonstrations of phase-matched and phase-mismatched filters aligned to an edge. Note that, while the phase-mismatched filter reduces contrast, it appears similarly visible to the phase-matched edge.

## Experiment 2

The results presented above reveal that the alignment of target filters with the spatial structure of the background determines detection sensitivity at least somewhat independently of target-background similarity. In Experiment 2 we tested our hypothesis that aligned targets are easier to detect because their amplitude is additive with the background amplitude, increasing local contrast. We therefore compared detection sensitivity to target filters that were either aligned or misaligned, and were either phase-matched or phase-mismatched with the original source image.

### Methods

All methods were identical to those of the preceding experiment, with the following changes. This experiment was carried out in our testing lab on a Display++ monitor (Cambridge Research Systems) with 14bit luminance precision (i.e. our local lockdown had lifted). The experimental design was a 2 (alignment: aligned versus misaligned) x 2 (phase: phase-matched versus phase-mismatched) factorial design (see Figure 8 for example stimuli). All target filters had an amplitude of 0.15, which, based on the data shown in Figure 4, we expected to yield a mean *d*’ of approximately 1. In all trials, there were four filters. Importantly, on half the trials, target filters were blended with the background as per Experiment 1, whereas in the other half of the trials the phase of the filters were reversed before blending. Note that, in the misaligned condition, the phase of the filters relative to the background is somewhat arbitrary, so in the analysis we average across these trials. Each observer completed a total of 800 trials, giving 200 trials per unique condition demonstrated in Figure 8. Testing took approximately 30 minutes per observer.

### Results

In Experiment 2, we included conditions that provided an opportunity to replicate our findings from Experiment 1. As demonstrated in the top left panel of Figure 8, we included a condition in which target filters were both aligned and phase-matched to the natural background structure, as per the aligned condition of Experiment 1. We first compare observers’ accuracy in this condition with the accuracy in the misaligned condition. The results are shown as connected points in Figure 9A, and reveal better performance in the (phase-matched) aligned condition than the misaligned condition for all observers. We therefore replicate the results from Experiment 1 under strict laboratory conditions. Also shown in Figure 9A are the results from the phase-mismatched condition, in which target filters were aligned with their backgrounds, but had their phase inverted. Importantly, phase-matched and phase-mismatched targets were well equated on similarity (phase-matched and phase-mismatched average similarities were .7 and 0.71, respectively). In the phase-mismatched condition, however, all observers were close to chance level, revealing they were unable to detect these targets (mean accuracy = 49%; RR = 45%, RW = 50%, WH = 51%).

**Figure 9.**
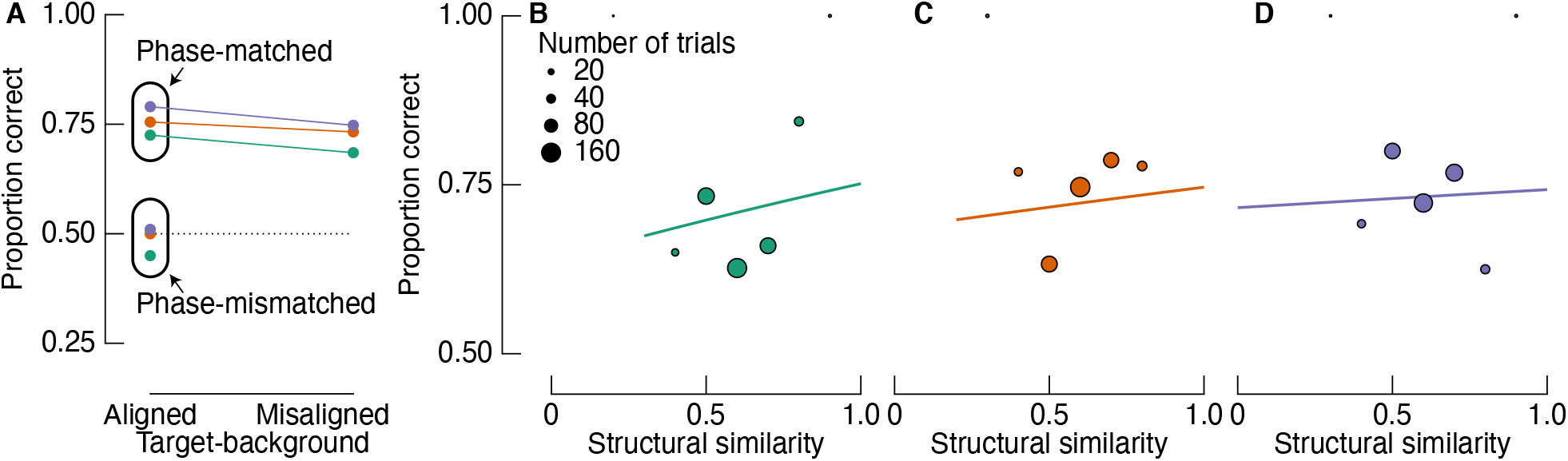
Proportion of correct target detections in Experiment 2. A) Results are shown according to the two main experimental factors: target-background alignment (x-axis) and target-background phase alignment (grouped data points). Note that target-background similarity is matched across aligned conditions (see left column of Figure 8). Colours indicate different observers as per Figure 3. B-D) Proportion correct as a function of target-background similarity for misaligned trials only. Solid lines are fits from a logistic GLMM, showing a positive relationship between similarity and detection accuracy, in contrast to the fits of Experiment 1 data and results reported by Sebastian et al (2017).

We again tested whether there was an inverse relationship between accuracy and similarity in the condition most closely matching the condition tested by Sebastian et al (2017), i.e. the misaligned condition (see Figure 7). We again used a logistic GLMM, and similarity was binned in 0.1 steps. In contrast to the fitted model in Experiment 1, however, we found a non-significant positive relationship between similarity and proportion correct (slope = 0.33, population standard deviation = 0.28, p = 0.596). Fits to observers’ data are shown in Figure 9B-D.

### Discussion

The aim of Experiment 2 was to test the prediction that targets aligned with their natural image backgrounds are easier to identify than targets that are misaligned with their backgrounds (i.e. the results of Experiment 1) due to a difference in local contrast. The differences in local contrast across these conditions results from contrast additivity in the aligned condition when filters are phase-matched with their background. We tested this prediction in Experiment 2 by inverting the phase of target filters on half of the trials in which the filters were aligned with the background structure. Inverting the phase of target filters aligned with their background has a subtractive effect on local contrast, and, as we expected, rendered observers incapable of detecting the targets (Figure 9). The results of Experiment 2 thus support the notion that target-background similarity is not a useful metric of the detectability of targets per se, whereas the interaction between relative phase and structural alignment is critical.

### Generative model of task performance

We next aimed to develop a model that captures the key results reported for Experiment 1 and Experiment 2. We hoped to account for the finding that aligned targets are more accurately detected than misaligned targets, and that this effect of alignment depends on the relative phase of target and background. These effects suggest that observers are tuning to local changes in the images caused by the additivity of filter and background luminance. The model is therefore based on simple luminance and contrast detection mechanisms like those involved in the generation of our stimuli (Figure 1). On each trial, the model detects difference in various image statistics across the target and distractor images, and then generates a response based on these differences. Simulated responses were determined by fitting the model output to observers’ responses in Experiment 1. We then tested whether the fitted model reproduced the key Experiment 2 results.

On each trial, the model was given the target image and the distractor image. The maximum contrast of each image was found by taking the maximum value of contrast maps as computed in Equations 1 – 6. The maximum luminance extreme of each image was the maximum absolute deviation of each image from mid-grey, capturing both local minima and maxima in images across trials. For each trial and each metric, we computed a ratio between left and right images:

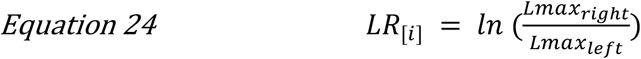

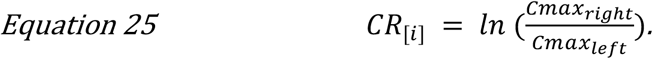

LR and CR refer to ratios of the most extreme luminance and contrast values, respectively, where negative values indicate greater extremes in the left image, and positive values indicate greater extremes in the right image. We weighted these metrics by fitting them to observers’ responses (i.e. left or right spatial interval) using logistic regression. We then analysed the model predictions as per the behavioural analyses. We built different models that included 1) just the absolute luminance peak of stimuli, 2) just the contrast energy maxima of stimuli, or 3) both.

The best model was one that detects the absolute luminance peak *and* the maximum contrast energy within the target and distractor images. As shown in Figure 10, this model reproduces the qualitative patterns of performance observed in Experiment 1 (compare the model data in Figure 10 with the empirical data in Figure 3). Importantly, each image statistic significantly contributes to the model (*p*’s < 0.001), and including both parameters provided a better fit than including either parameter alone based on formal model comparison (chi-square test compared with the next best model: *χ*^2^(Δ*df* = 4) = 114.3, *p* < 0.001). Note that the generative model responses are fit to the observer data based solely on the image-computable features, not on the labels of the experimental conditions (e.g. the data are not fit to aligned vs misaligned conditions) – yet the model reproduces the same patterns of data across conditions as observers. Adding the model’s predicted response to the signal detection model also improved estimates of observers’ *d*’(*χ*^2^(Δ*df* = 6) = 163.1, *p* < 0.001). These model results are consistent with the notion that adding filters to the image causes local peaks in absolute luminance and contrast that can differentiate the target image from the distractor image (Bex & Makous, 2002). As described next, however, this model cannot account for the findings in Experiment 2.

**Figure 10.**
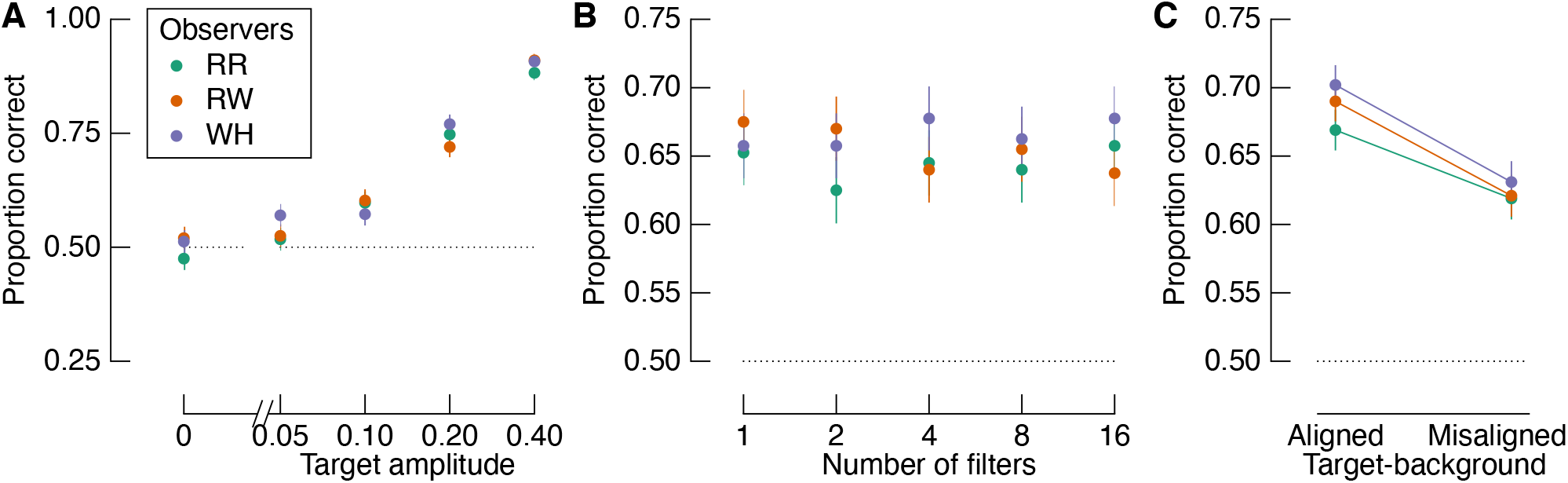
Model performance for Experiment 1. Compare with Figure 3. The model captures the key results from Experiment 1. Error bars show one binomial standard deviation.

We fit the model to observers’ responses from Experiment 1, as shown above, and tested whether the fitted model could predict observers’ responses to Experiment 2. The critical test is whether the model falls to chance when filters were spatially aligned but phase-inverted relative to their backgrounds (i.e. the phase-mismatched condition). As shown in Figure 11, however, model performance was well *below* chance in this condition. This below-chance performance occurs because phase-inversion reduces the luminance and contrast peaks of the target image to below those of the distractor image, resulting in the model reporting the distractor as the target more often than not. The model again reproduces the effect of alignment, but overestimates the size of the effect. The overestimation may have resulted from the changes we made between experiments, including using different displays and filter amplitudes. These differences are less relevant than the model’s gross misestimation of the phase-mismatched condition as described below.

**Figure 11.**
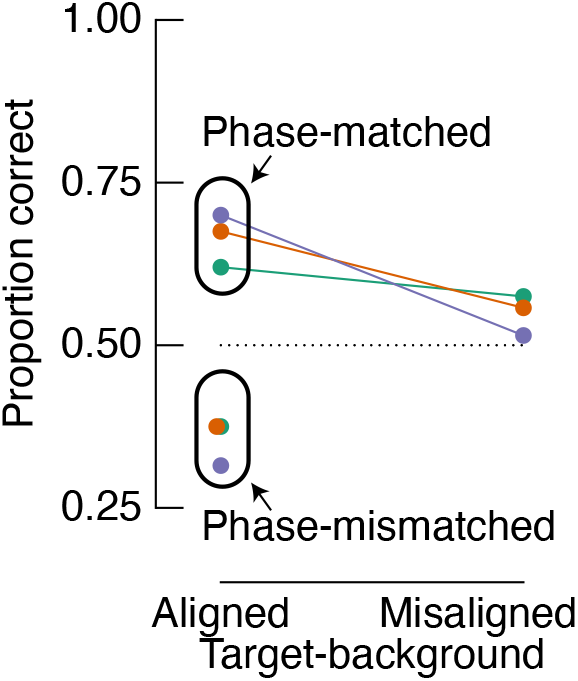
Model performance for Experiment 2. Compare with Figure 9A. The model performs below chance in the phase mismatched condition, whereas observers were at chance (Figure 9A).

The relatively poor fit to the phase-mismatched condition of Experiment 2 is informative. Observers’ data cannot be explained fully by assuming that they adopted a simple rule in which they detected luminance and contrast peaks. For phase-matched trials, therefore, the superior sensitivity to aligned filters over misaligned filters cannot be solely accounted for by tuning to local peaks. We attempted to improve the fit using other image metrics, such as contrast energy combined across spatial scales and the energy of a “back pocket” filter-rectify-filter model (Harrison & Bex, 2016; Landy, 2013), but none improved the fits of the model. Rather than taking the maximum contrast or luminance extreme, we also tried using k-maxima (up to k = 1000), which also did not improve the fits. Observers’ performance, therefore, escapes a relatively straightforward low-level explanation.

## General Discussion

The aim of the present study was to test observers’ ability to detect contrast-defined targets that have been blended with natural image backgrounds. We designed the target filters so that their orientations were either aligned or misaligned with the local background structure. Based on the recent report that detection sensitivity is inversely related to target-background similarity (Sebastian et al., 2017), we expected to find worse performance in the aligned condition relative to the misaligned condition based on the notion that aligned targets would have higher target-background similarity than misaligned targets. Across two experiments, however, we found superior detection of targets that were designed to be aligned and phased-matched to the structure of the background compared with targets that were designed to be misaligned with their backgrounds and were thus lower in similarity. As noted below, our goal was not to replicate the study by Sebastian et al, but instead to test the role of target-background similarity in target detection using a novel approach.

Our experiments show that observers’ sensitivity does not linearly scale with similarity in all cases. In Experiment 1 we found a positive relationship between similarity and sensitivity: observers were most sensitive to targets in which target filters were aligned with, and most similar to, background structure (Figure 3C and Figure 4C). We replicated this finding in Experiment 2 (Figure 9A). When we limited our analyses to only trials in which target filters were misaligned with the background structure, we found mixed results across the two experiments: a negative relationship between detection accuracy and target-background similarity in Experiment 1 (Figure 7), but a positive relationship in Experiment 2 (Figure 9C-D). The cause of this difference in results across experiments is not clear, but we note that we did not design either experiment to specifically measure the relationship between similarity and sensitivity in this way, and neither model was significant. Regardless, there was no clear evidence of an inverse relationship as we expected.

The limitation of a specific structural similarity metric as a predictor of sensitivity in our study is most apparent in our Experiment 2 results. By matching or inverting the phase of filters aligned with the background structure, we produced target-background images that were equivalent in similarity but were different in their detectability (Figure 8 and Figure 9A). When phase was inverted relative to the background, observers’ performance was at chance level. The phase-mismatched condition therefore removed the information observers depended on to perform the task (i.e. contrast). A variant of the metric of similarity used here and by Sebastian et al (2017) may better predict sensitivity if it encodes phase information. Computationally, similarity is analogous to a normalised correlation coefficient; retaining phase would yield similarity scores ranging from −1 (perfectly matched counter-phase) and 1 (perfectly matched in-phase). However, Sebastian et al. used targets and a template-matching ideal observer model that had a fixed phase, in which case sensitivity may indeed scale inversely with phase-invariant similarity. We also note that the similarity between target filters and backgrounds in our study was approximately double those reported by Sebastian et al, and so it is possible that a linear inverse influence of similarity on sensitivity holds for relatively low levels of similarity.

In Experiment 1, there was no clear relationship between the number of filters added to the target image and observers’ performance (Figure 3B and Figure 4B). We expected to find such a relationship based on the simple principle that there is an additional opportunity to detect a target for each filter added (Macmillan & Creelman, 2004). On closer inspection of the proportion correct data presented in Figure 3B, only a single data point in each of RW and WH’s data are inconsistent with the expected trend for all observers. The lack of a statistically robust finding, therefore, may be due to our limited number of observers. A clear main effect of the number of filters was possibly also obscured by an interaction with target amplitude as shown in Figure 5. It is interesting that our generative model also produced a somewhat noisy relationship between accuracy and the number of filters. It is possible that observers outperformed our model in the phase-mismatched condition of Experiment 2 because they integrated information over multiple locations rather than using the maximum in each image. However, even when our model had access to the top 1000 maxima in the images, it still performed below chance.

Detection thresholds in our experiments are approximately an order of magnitude greater than those reported by Sebastian et al. This is not particularly surprising given that we did not attempt to replicate their design, and instead used stimuli and methods that differed from theirs. One aspect of our experiments that would have likely decreased sensitivity was the lack of spatial certainty in the position of targets. Target filters could appear anywhere within the natural image background, maximising spatial uncertainty. The ability to identify targets depends on spatial (un)certainty, particularly in peripheral vision (Bennett & Banks, 1991; Harrison & Bex, 2015, 2017; Levi et al., 1987). Sebastian et al reduced spatial uncertainty by presenting targets at the same location on each target-present trial. Lower thresholds should be expected with such reduced uncertainty relative to our experiment in which observers had to search the entire background region. When observers are required to search for a potential target in a new background, false alarms can occur anywhere in the image. Computationally, such search can be performed using the same basic processes as involved in detection of a target at a cued location. In addition to determining whether a filter response is greater than a threshold (e.g. Sebastian et al., 2017), however, search involves determining which of several locations is most likely a target. We modelled this by taking the spatial interval with maximum luminance and contrast energy.

Despite the differences between our study and previous studies, we can confidently conclude that phase plays an important role in target detection for at least the sorts of targets used in our study (i.e. directional first-derivative of Gaussians). This result was foreshadowed by Bex and Makous (2002), who suggested detection thresholds for natural images depend on local phase-alignment within or across frequency bands. Our modelling suggests that observers detected the target interval by using local luminance extremes and contrast maxima. These local visual cues were most apparent in conditions in which the target phase was additive with the background. Phase, therefore, played an important role in our experiments. However, the failure of this model to capture observers’ accuracy in the phase-mismatched condition of Experiment 2 reveals that a rule using local extremes is overly simple. We do not think it is likely that observers were switching strategies across conditions, because observers could not have known on a given trial which condition was displayed. In phase-mismatched trials in which the extremes of the target image were reduced to below the level of the extremes of the distractor image, observers must have been using other image cues that have escaped our description. Anecdotally, all observers, who are experienced psychophysical observers, reported using a template-matching strategy. This insight is obviously limited in its usefulness, because a template-matching strategy is equivalent to the computations performed in our model.

Our results are consistent with those of Neri (2011), who investigated the influence of target phase relative to the structure of a natural image background and how these effects differ when the stimulus is inverted. Neri found that, when the background is upright, observers tune to feature detectors that are in-phase with a natural edge. When the background is inverted, observers’ tuning is less phase-aligned with the natural edge. These results suggest that upright scenes produce a bias in visual processing that steers observers’ filtering toward the local phase of natural structures. These results are consistent with our own finding that observers are most sensitive to filters that are spatially aligned, and in-phase, with the background.

The extent to which the design of visual targets determines detectability in natural backgrounds is thus clearly an important consideration. As shown in a demonstration by Sebastian et al, it is incontrovertible that there are some targets for which phase is unimportant for visibility. We reproduce such a demonstration in the top row of Figure 12. We question, however, the relevance of a similarity metric in explaining the visibility of the target in this demonstration, considering that similar demonstrations can be produced in which target-background similarity is greater, and yet the target is easily visible (bottom row, Figure 12). The importance of target-background phase (in)variance likely depends on multiple factors, such as target design, as well as differences in the background in the region of the target (i.e. ‘partial masking’, see Sebastian et al., 2020). In addition to testing target visibility in different backgrounds based on contrast, luminance, and similarity (Sebastian et al., 2017, 2020), binning backgrounds according to their phase-similarity with targets may clarify these interactions in future experiments.

**Figure 12.**
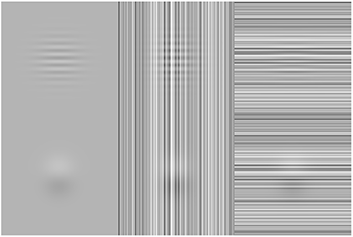
Similarity alone is a poor predictor of visibility. Top and bottom panels show a 16 cyc/image target and Gaussian-derivative target, respectively, in each of three backgrounds (left to right): zero noise, vertical noise, and horizontal noise. Noise is 1D Gaussian (standard deviation = 0.15), and targets have the same amplitude in all panels (0.1). Similarity masking is demonstrated in the top right panel, in which the high-frequency target is rendered invisible. Such a masking effect is phase-invariant. As shown in the bottom right panel, however, a broader-band target with the same amplitude remains unmasked by the same noise. Importantly, the structural similarity between the target and background is greatest in the bottom right panel (0.15 bottom right vs 0.12 in the top right).

We used images from the THINGS database as naturalistic backgrounds. The THINGS database is a recently released database with over 26,000 images from 1854 categories (Hebart et al., 2019). Relatively little has been reported about the basic statistical properties of the images in this database, and so it is possible that they may deviate from what one may expect from typical natural images ^1^. In Figure 13 we show the mean image spectra as a function of spatial frequency and orientation. This analysis shows that THINGS images have, on average, the same basic image spectra as typically found in natural images: contrast energy decreases with increasing spatial frequency (Figure 13A), and there is an over representation of cardinal orientations (Figure 13B). It is therefore unlikely that something in particular about the distribution of contrast energy in the THINGS images played a key role in our results. However, it is likely that the visual system performs different operations when processing visual objects like those in the THINGS images compared with visual textures (Wallis et al., 2019). Sebastian et al (2017) used images of scenes that had no particular focus on objects per se. To the best of our knowledge, no study has systematically investigated whether target detection differs on backgrounds of things versus backgrounds of stuff.

**Figure 13.**
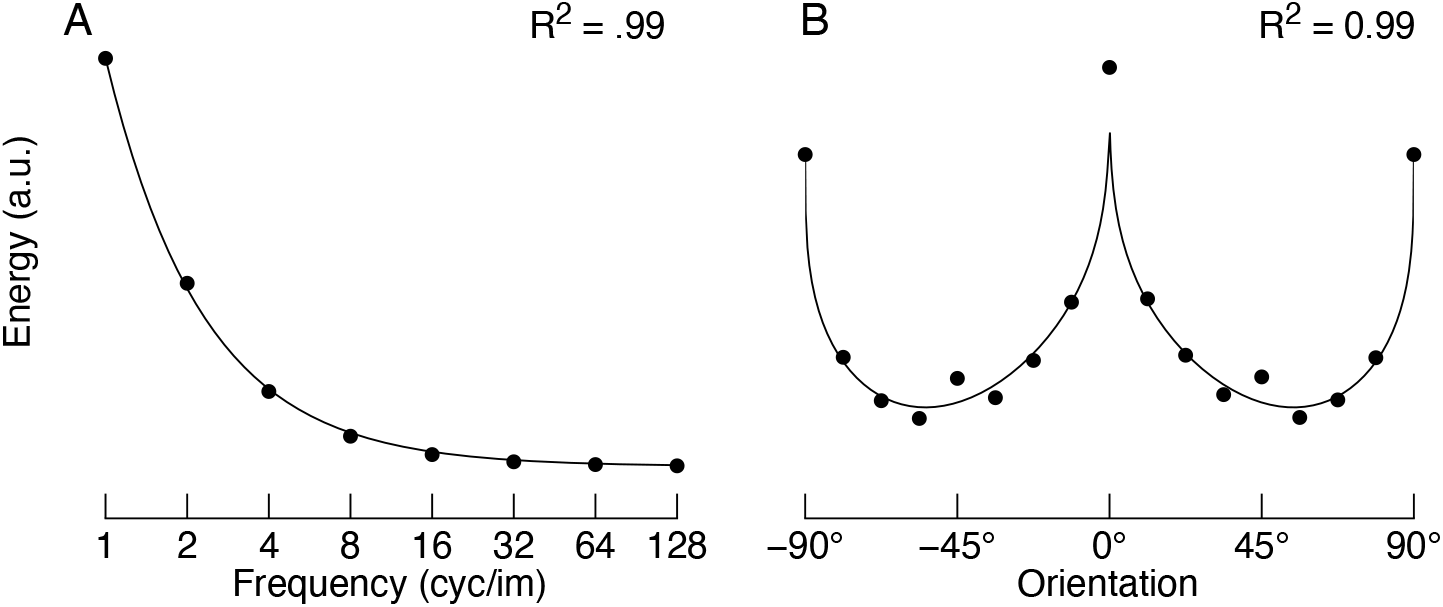
Mean spectra of all images in the THINGS database. A) Mean contrast energy as a function of spatial frequency. The solid line is the fit of the function 1/f^a^, which explains 99% of the variance. The free parameter, a, was 1.2. B) Mean contrast energy as a function of orientation. We computed contrast energy in the frequency domain using a series of raised cosine filters centred on a given orientation and spanning all spatial frequencies (“bow-tie filters”; full width half height = 12.5°). These data were then fit with a function that captures the over-representation of cardinals, as well as the greater contrast energy for horizontal contrast compared with vertical contrast ^2^. This function explains 99% of the variance.

Several studies suggest that there are high-level influences over the tuning of low-level feature detectors like those used to detect targets in the present study. For example, Teufel et al (2018) found that prior knowledge about image content influences the detectability of oriented targets aligned to locally occluded edges. Neri (2017) similarly found that sensitivity is greatest on edges implied by image content, regardless of whether local contrast detectors would respond at the region of the target. Harrison & Rideaux (2019) further showed that edge detection in visual noise can be greatly influenced by the allocation of visual attention. Taken together, these findings suggest that the detectability of targets in the present study may have depended on the specific objects in background images, and how combinations of low-level and high-level factors guided observers’ visual attention. However, we did not design our experiments to examine such possible differences across object images. The availability of repositories such as the THINGS database makes such questions possible to address in future studies.

In summary, we tested observers’ ability to detect targets in natural image backgrounds. Observers were most sensitive to targets when they were aligned and phase-matched with their backgrounds. Inverting the phase of aligned targets reduced observers’ detection performance to chance. To best model the image factors that predict human sensitivity to contrast defined targets in natural backgrounds, therefore, the phase of the target relative to the background must be considered.

## Author contributions

WJH and PJB designed the stimulus generation method and conceived the general idea. WJH designed Experiment 1. WJH, RR, and JBM designed Experiment 2. WJH, RR, and RKW conducted the experiments. WJH conducted all analyses and generated all figures. TSAW advised on the multilevel modelling and was involved in many discussions leading to the conception of the study. RR, RKW and WJH wrote the first draft of the manuscript. All authors reviewed and edited the manuscript.

## Acknowledgements

This research was supported by an Australian Research Council Discovery Early Career Award to WJH (DE190100136) and RR (DE210100790). JBM was supported by the Australian Research Council (ARC) Centre of Excellence for Integrative Brain Function (ARC Centre Grant CE140100007). PJB was supported by a National Institute of Health R01 (EY029713).

## Footnotes

1 Since originally submitting this manuscript, a more thorough description of the luminance and luminance contrast properties of THINGS images was presented by Harrison (2021).

2 This function has the form: *E* = (*a* – *mag* * |*sin*(2*θ*)|^b^) * (*p* * (*cos*(2*θ*)) + *p* + 1), where E is contrast energy, *θ* is orientation in radians, and *a, mag, b*, and *p* are free parameters. See Harrison (2021) for a more detailed analysis of the THINGS images.

## References

Balas, B., Nakano, L., & Rosenholtz, R. (2009). A summary-statistic representation in peripheral vision explains visual crowding. Journal of Vision, 9(12), 1–18. https://doi.org/10.1167/9.12.13

Barlow, H. B. (1961). Possible Principles Underlying the Transformations of Sensory Messages. In W. A. Rosenblith (Ed.), Sensory Communication (Vol. 1, pp. 216–234). The MIT Press. https://doi.org/10.7551/mitpress/9780262518420.003.0013

Barlow, H. B. (1972). Single Units and Sensation: A Neuron Doctrine for Perceptual Psychology? Perception, 1(4), 371–394. https://doi.org/10.1068/p010371

Bates, D., Mächler, M., Bolker, B., & Walker, S. (2015). Fitting Linear Mixed-Effects Models Using lme4. Journal of Statistical Software, 67(1), 1–48. https://doi.org/10.18637/jss.v067.i01

Bennett, P. J., & Banks, M. S. (1991). The effects of contrast, spatial scale, and orientation on foveal and peripheral phase discrimination. Vision Research, 31(10), 1759–1786.

Bex, P. J., & Makous, W. (2002). Spatial frequency, phase, and the contrast of natural images. Journal of the Optical Society of America A, 19(6), 1096–1106. https://doi.org/10.1364/JOSAA.19.001096

Bex, P. J., Solomon, S. G., & Dakin, S. C. (2009). Contrast sensitivity in natural scenes depends on edge as well as spatial frequency structure. Journal of Vision, 9(10), 1–19. https://doi.org/10.1167/9.10.1

Brainard, D. H. (1997). The Psychophysics Toolbox. Spatial Vision, 10(4), 433–436.

Cadena, S. A., Denfield, G. H., Walker, E. Y., Gatys, L. A., Tolias, A. S., Bethge, M., & Ecker, A. S. (2019). Deep convolutional models improve predictions of macaque V1 responses to natural images. PLOS Computational Biology, 15(4), e1006897. https://doi.org/10.1371/journal.pcbi.1006897

Campbell, F. W., & Robson, J. G. (1968). Application of Fourier analysis to the visibility of gratings. The Journal of Physiology, 197(3), 551–566.

Carandini, M., Demb, J. B., Mante, V., Tolhurst, D. J., Dan, Y., Olshausen, B. A., Gallant, J. L., & Rust, N. C. (2005). Do we know what the early visual system does? Journal of Neuroscience, 25(46), 10577–10597. https://doi.org/10.1523/JNEUROSCI.3726-05.2005

Dorr, M., & Bex, P. J. (2013). Peri-Saccadic Natural Vision. Journal of Neuroscience, 33(3), 1211–1217. https://doi.org/10.1523/JNEUROSCI.4344-12.2013

Field, D. J. (1987). Relations between the statistics of natural images and the response properties of cortical cells. JOSA A, 4(12), 2379–2394. https://doi.org/10.1364/JOSAA.4.002379

Freeman, W. T., & Adelson, E. H. (1991). The design and use of steerable filters. IEEE Transactions on Pattern Analysis &\ldots. http://www.computer.org/csdl/trans/tp/1991/09/i0891.pdf

Geisler, W. S. (2008). Visual Perception and the Statistical Properties of Natural Scenes. Annual Review of Psychology, 59(1), 167–192. https://doi.org/10.1146/annurev.psych.58.110405.085632

Gelman, A., & Hill, J. (2007). Data Analysis Using Regression and Multilevel/Hierarchical Models. Cambridge University Press.

Greenwood, J. A., Bex, P. J., & Dakin, S. C. (2010). Crowding changes appearance. Current Biology, 20(6), 496–501. https://doi.org/10.1016/j.cub.2010.01.023

Greenwood, J. A., Bex, P. J., & Dakin, S. C. (2012). Crowding follows the binding of relative position and orientation. Journal of Vision, 12(3). https://doi.org/10.1167/12.3.18

Harrison, W. J. (2021). Luminance and contrast of images in the THINGS database. BioRxiv, 2021.07.08.451706. https://doi.org/10.1101/2021.07.08.451706

Harrison, W. J., & Bex, P. J. (2014). Integrating retinotopic features in spatiotopic coordinates. Journal of Neuroscience, 34(21), 7351–7360. https://doi.org/10.1523/JNEUROSCI.5252-13.2014

Harrison, W. J., & Bex, P. J. (2015). A unifying model of orientation crowding in peripheral vision. Current Biology, 25(24), 3213–3219. https://doi.org/10.1016/j.cub.2015.10.052

Harrison, W. J., & Bex, P. J. (2016). Reply to Pachai et al. Current Biology, 26(9), R353–R354. https://doi.org/10.1016/j.cub.2016.03.024

Harrison, W. J., & Bex, P. J. (2017). Visual crowding is a combination of an increase of positional uncertainty, source confusion, and featural averaging. Scientific Reports, 7, 45551. https://doi.org/10.1038/srep45551

Harrison, W. J., & Rideaux, R. (2019). Voluntary control of illusory contour formation. Attention, Perception, and Psychophysics, 81(5), 1522–1531. https://doi.org/10.3758/s13414-019-01678-8

Haun, A. M., & Peli, E. (2013). Perceived contrast in complex images. Journal of Vision, 13(13), 3. https://doi.org/10.1167/13.13.3

Hebart, M. N., Dickter, A. H., Kidder, A., Kwok, W. Y., Corriveau, A., Wicklin, C. V., & Baker, C. I. (2019). THINGS: A database of 1,854 object concepts and more than 26,000 naturalistic object images. PLOS ONE, 14(10), e0223792. https://doi.org/10.1371/journal.pone.0223792

Hubel, D. H., & Wiesel, T. N. (1959). Receptive fields of single neurones in the cat’s striate cortex. The Journal of Physiology, 148, 574–591.

Kleiner, M., Brainard, D., Pelli, D., Ingling, A., Murray, R., & Broussard, C. (2007). What’s new in psychtoolbox-3. Perception, 36(14), 1–16.

Landy, M. S. (2013). Texture analysis and perception. In The New Visual Neurosciences (pp. 639–652). MIT Press.

Levi, D. M., Klein, S. A., & Yap, Y. L. (1987). Positional uncertainty in peripheral and amblyopic vision. Vision Research, 27(4), 581–597.

Macmillan, N. A., & Creelman, C. D. (2004). Detection theory: A user’s guide. https://doi.org/10.1016/0376-6357(93)90083-4

Neri, P. (2011). Global Properties of Natural Scenes Shape Local Properties of Human Edge Detectors. Frontiers in Psychology, 2. https://doi.org/10.3389/fpsyg.2011.00172

Neri, P. (2014). Dynamic engagement of human motion detectors across space-time coordinates. Journal of Neuroscience, 34(25), 8449–8461. https://doi.org/10.1523/JNEUROSCI.5434-13.2014

Neri, P. (2017). Object segmentation controls image reconstruction from natural scenes. Plos Biology, 15(8), e1002611. https://doi.org/10.1371/journal.pbio.1002611

Olshausen, B. A., & Field, D. J. (2005). How close are we to understanding v1? Neural Computation, 17(8), 1665–1699. https://doi.org/10.1162/0899766054026639

Parraga, C. A., Troscianko, T., & Tolhurst, D. J. (2000). The human visual system is optimised for processing the spatial information in natural visual images. Current Biology. http://www.sciencedirect.com/science/article/pii/S0960982299002626

Pelli, D. G. (1997). The VideoToolbox software for visual psychophysics: Transforming numbers into movies. Spatial Vision, 10(4), 437–442.

Rosenholtz, R., Huang, J., & Ehinger, K. A. (2012). Rethinking the role of top-down attention in vision: Effects attributable to a lossy representation in peripheral vision. Frontiers in Psychology, 3, 13. https://doi.org/10.3389/fpsyg.2012.00013

Sebastian, S., Abrams, J., & Geisler, W. S. (2017). Constrained sampling experiments reveal principles of detection in natural scenes. Proceedings of the National Academy of Sciences of the United States of America, 114(28), E5731–E5740. https://doi.org/10.1073/pnas.1619487114

Sebastian, S., Seemiller, E. S., & Geisler, W. S. (2020). Local reliability weighting explains identification of partially masked objects in natural images. Proceedings of the National Academy of Sciences, 117(47), 29363–29370. https://doi.org/10.1073/pnas.1912331117

Simoncelli, E. P., & Olshausen, B. A. (2001). Natural image statistics and neural representation. Annual Review of Neuroscience, 24, 1193–1216. https://doi.org/10.1146/annurev.neuro.24.1.1193

Smith, P. L., & Little, D. R. (2018). Small is beautiful: In defense of the small-N design. Psychonomic Bulletin & Review, 1–19. https://doi.org/10.3758/s13423-018-1451-8

Teufel, C., Dakin, S. C., & Fletcher, P. C. (2018). Prior object-knowledge sharpens properties of early visual feature-detectors. Scientific Reports, 8(1), 10853. https://doi.org/10.1038/s41598-018-28845-5

Wallis, T. S. A., & Bex, P. J. (2012). Image correlates of crowding in natural scenes. Journal of Vision, 12(7), 1–19. https://doi.org/10.1167/12.7.6

Wallis, T. S. A., Dorr, M., & Bex, P. J. (2015). Sensitivity to gaze-contingent contrast increments in naturalistic movies: An exploratory report and model comparison. Journal of Vision, 15(8), 3. https://doi.org/10.1167/15.8.3

Wallis, T. S. A., Funke, C. M., Ecker, A. S., Gatys, L. A., Wichmann, F. A., & Bethge, M. (2019). Image content is more important than Bouma’s Law for scene metamers. ELife, 8, e42512. https://doi.org/10.7554/eLife.42512

